# Architect: a tool for producing high-quality metabolic models through improved enzyme annotation

**DOI:** 10.1101/2021.10.12.464133

**Authors:** Nirvana Nursimulu, Alan M. Moses, John Parkinson

## Abstract

**Motivation:** Constraints-based modeling is a powerful framework for understanding growth of organisms. Results from such simulation experiments can be affected at least in part by the quality of the metabolic models used. Reconstructing a metabolic network manually can produce a high-quality metabolic model but is a time-consuming task. At the same time, current methods for automating the process typically transfer metabolic function based on sequence similarity, a process known to produce many false positives.

**Results:** We created Architect, a pipeline for automatic metabolic model reconstruction from protein sequences. First, it performs enzyme annotation through an ensemble approach, whereby a likelihood score is computed for an EC prediction based on predictions from existing tools; for this step, our method shows both increased precision and recall compared to individual tools. Next, Architect uses these annotations to construct a high-quality metabolic network which is then gap-filled based on likelihood scores from the ensemble approach. The resulting metabolic model is output in SBML format, suitable for constraints-based analyses. Through comparisons of enzyme annotations and curated metabolic models, we demonstrate improved performance of Architect over other state-of-the-art tools.

**Availability:** Code for Architect is available at https://github.com/ParkinsonLab/Architect.

**Supplementary information:** Supplementary data are available at *Bioinformatics* online.

## 1. INTRODUCTION

Metabolic modeling has been used for engineering strains of bacteria for bioremediation, for understanding what drives parasite growth, as well as for shedding light on how disruptions in the microbiome can lead to progression of various diseases (Bauer & Thiele, 2018; Nemr et al., 2018; Song et al., 2013). In any of these applications, the standard protocol is to first construct an initial draft of the metabolic model of the organism(s) (consisting of the biochemical reactions predicted present) followed by a gap-filling procedure, whereby additional reactions are introduced to ensure that simulations can be performed (Pan & Reed, 2018). Importantly, errors introduced at any steps of model reconstruction can impact downstream simulations and result interpretation (Thiele & Palsson, 2010). For instance, false positive enzyme predictions may mask the essentiality of key pathways; on the other hand, the organism’s metabolic abilities may be underestimated when metabolic enzymes and pathways are incorrectly left out or under-predicted (Guzman et al., 2015). While these concerns can be addressed through dedicated manual curation, such efforts tend to be extremely time-consuming. Instead, attention has turned to the use of automated methods, such as PRIAM and CarveMe, the latter capable of generating fully functional genome-scale metabolic models (Claudel-Renard, Chevalet, Faraut, & Kahn, 2003; Machado, Andrejev, Tramontano, & Patil, 2018). Given a genome of interest, CarveMe uses sequence similarity searches to assign confidence scores to reactions within a universal model of metabolism. Based on these scores, a genome-specific metabolic model is then reconstructed by removing reactions that are either not identified or poorly supported, and adding in reactions to fill gaps to construct functional pathways (Machado et al., 2018).

A key step in this process is the accurate identification of enzymes based on sequence data alone and can be formally defined as follows: given an amino acid sequence, what are its associated enzymatic function(s), if any? The problem is a multi-label classification problem; here we consider enzymatic functions as defined by the Enzyme Commission (EC), in which enzymes are assigned to EC numbers representing a top-down hierarchy of function (Bairoch, 2000). Enzyme annotation can be performed by inferring homology to known enzymes based on sequence similarity searches using methods such as BLAST and DIAMOND (Altschul, Gish, Miller, Myers, & Lipman, 1990; Buchfink, Xie, & Huson, 2015). However, such methods do not consider the overlap of sequence similarity between enzyme classes and are prone to an unacceptable rate of false positive predictions (Hung, Wasmuth, Sanford, & Parkinson, 2010). To overcome such errors, a number of more specialized tools have been developed that take advantage of sequence features or profiles specific to individual enzyme classes (Claudel-Renard et al., 2003; Hung et al., 2010; Nguyen, Srihari, Leong, & Chong, 2015; Nursimulu, Xu, Wasmuth, Krukov, & Parkinson, 2018). For example, DETECT (Density Estimation Tool for Enzyme ClassificaTion) considers the effect of sequence diversity when predicting different enzyme classes (Hung et al., 2010; Nursimulu et al., 2018), while PRIAM and EnzDP rely on searches of sequence profiles constructed from families of enzymes (Claudel-Renard et al., 2003; Nguyen et al., 2015). Each tool provides different advantages in terms of accuracy and coverage.

Here we present Architect, a tool capable of automatically constructing a functional metabolic model for an organism of interest, based on its proteome. At its core, Architect exploits an ensemble approach that combines the unique strengths of multiple enzyme annotation tools to ensure high confidence enzyme predictions. Subsequently gap-filling is performed to construct a functional metabolic model in Systems Biology Markup Language (SBML) format which can be readily analysed by existing constraints-based modeling software. In parallel with its namesake, Architect not only designs the metabolic model of an organism, but it also coordinates the sequence of steps that go towards the SBML output given user specifications, such as the definition of an objective function for gap-filling. We evaluate the performance of Architect both in terms of its ability to perform accurate enzyme annotations, relying on UniProt/SwissProt sequences as a gold standard (The UniProt Consortium, 2014) and, separately, as a metabolic model reconstruction tool by focusing on 3 organisms for which curated metabolic models have already been generated (*Caenorhabditis elegans* (Witting et al., 2018), *Neisseria meningitidis* (Mendum, Newcombe, Mannan, Kierzek, & McFadden, 2011) and *E. coli* (Monk et al., 2017)). Compared to other state-of-the-art methods (Claudel-Renard et al., 2003; Machado et al., 2018), Architect delivers improved performance both in terms of enzyme annotation and model reconstruction.

## 2. METHODS

### 2.1 Sources of Sequence Data

Sequences were downloaded from the SwissProt database (The UniProt Consortium, 2014) and their corresponding annotations from the ENZYME database (Bairoch, 2000) (downloaded on February 9^th^, 2021). Only complete EC numbers were considered in building Architect’s ensemble classifiers. Further, ECs associated with fewer than 10 protein sequences were removed to ensure sufficient training data. This filtering resulted in a final collection of 1,670 ECs represented by 207,121 sequences (**Supplemental Figure 1**). A further set of 294,067 protein sequences not associated with either complete or partial EC annotations (subsequently referred to as “non-enzymes”) was retrieved from the same version of SwissProt. For generation of test and training datasets for use in five-fold cross-validation steps, ECs associated with multifunctional proteins, were divided into appropriate sets using a previously published protocol (Sechidis, Tsoumakas, & Vlahavas, 2011).

### 2.2 Enzyme Annotation Using Ensemble Classifiers

For any given protein, EC predictions are generated through integrating the output from five state-of-the-art enzyme annotation tools: EFICAz v2.5.1 (Kumar & Skolnick, 2012), PRIAM version 2018 (Claudel-Renard et al., 2003), DETECT v2 (Nursimulu et al., 2018), EnzDP (Nguyen et al., 2015) and CatFam (Yu, Zavaljevski, Desai, & Reifman, 2009). In addition to examining the performance of two relatively simple approaches, majority rule (in which we take the prediction supported by the most tools) and EC-specific best tool (in which we take the prediction from the tool which is found to perform best for a specific EC), we also investigated the performance of the following three classifiers: (1) naïve Bayes, (2) logistic regression and (3) random forest. For training each method, we first find, for each EC *x*, positive (proteins actually of class *x*) and negative examples (other proteins predicted by any tool to have activity *x*). For each protein *i*, a feature vector is then constructed consisting of the level of confidence in each tool’s prediction (based on the confidence score output by the tool); the associated binary label *y_i_* indicates whether the *i*^th^ protein has activity *x* and thus has value 1 if and only if the protein has activity *x*. For any EC predicted by all tools without false positives (that is without proteins of other classes misclassified with the EC), we apply a rule whereby we automatically assign the EC if made by any of these tools. Otherwise, those predictions made by an ensemble method with a likelihood score greater than 0.5 are considered to be of high-confidence.

For the naïve Bayes classifier trained on high-confidence predictions, given an EC predictable by *k* tools (1 ≤ *k* ≤ 5 depending on the number of tools that can predict the EC), each protein sequence is assigned a corresponding feature vector *F* of length *k*, where *F_i_* = 1 if the EC was predicted with high-confidence by the *i*^th^ tool, and *F_i_* = 0 otherwise. The posterior probability of a new protein *j* having EC *x* (the aforementioned “likelihood score”) is then given by the following equation, where each feature is assumed to follow a Bernouilli distribution:

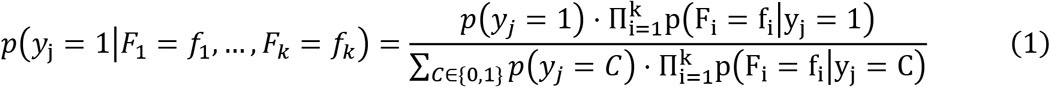

Other ensemble methods explicitly consider the level of confidence by each tool (see **Supplementary Material**). For example, our logistic regression classifiers train on feature vectors which use one-hot encoding to denote the level of confidence in the EC prediction by each tool. In the case of our random forest classifiers, each element of the feature vectors takes on a discrete value indicating the level of confidence by each tool.

Given that those ECs associated with fewer than 10 protein sequences are filtered out of the training data, some ECs may not be predictable by the classifier but may nevertheless be predicted by other tools; in particular, PRIAM consists of profiles specific to ECs associated with as few as a single sequence. To ensure higher coverage of metabolic reactions and pathways, EC predictions made with high-confidence by PRIAM are subsequently assigned as high-confidence during downstream model reconstruction.

### 2.3 Reconstruction of Metabolic Networks

From the set of high confidence enzyme annotations generated for the organism of interest (using the naïve Bayes classifier by default), an initial metabolic model is constructed with reference to either the Kyoto Encyclopedia of Genes and Genomes (KEGG) resource (Kanehisa, Furumichi, Sato, Ishiguro-Watanabe, & Tanabe, 2021) or the Biochemical, Genetic and Genomic (BiGG) knowledgebase (King et al., 2016). In brief, reaction identifiers and equations corresponding to high-confidence EC predictions are collated, along with non-enzymatic, spontaneous and any user-specified reactions (see **Supplementary Material**). Amongst the latter reactions, an objective function (such as biomass production) is required for downstream gap-filling. Furthermore, if BiGG is used as the reference reaction database, we also include non-EC associated reactions (including transport reactions). This step is performed through BLAST-based sequence similarity searches of the organism’s proteome against a database of protein sequences representing these non-EC associated reactions, collated from the BiGG resource, using a cut-off of 10^−20^ (Altschul et al., 1990; Buchfink et al., 2015).

Having generated an initial network, Architect next attempts to fill gaps within the network, representing reactions required to complete pathways necessary for the production of essential metabolites (as defined by the objective function). First a global set of candidate gap-filling reactions (*R*) is constructed by combining: 1) reactions that were previously identified in the enzyme annotation step at either low- or no-confidence; and 2) exchange reactions for dead-end metabolites (whose presence otherwise results in inactive (blocked) reactions that can inhibit biomass production (Ponce-de-León, Montero, & Peretó, 2013)). From this global set, Architect attempts to identify a set of reactions which, when supplemented to the initial network, is minimally sufficient for non-zero flux through the aforementioned user-defined objective function. This process leverages the mixed-integer linear programming (MILP) formulation employed by the CarveMe pipeline (Machado et al., 2018). First, penalties are assigned for the addition of each gap-filling candidate as follows. For the *i*^th^ reaction associated with 1 or more ECs predicted with low-confidence by the ensemble classifier (0.0001 < score ≤ 0.5), we find the highest score *S_i_* associated with any of the corresponding EC annotations. Then, we scale the scores of the gap-filling candidates to have a median of 1 (where *S_M_* is the median of the original scores):

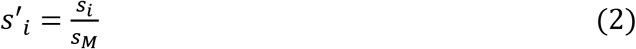

The penalty *p_i_* for adding the *i*^th^ reaction is then inversely proportional to the normalized score:

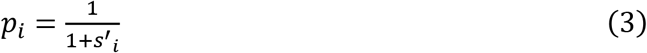

Remaining candidate reactions for gap-filling (that is, those either not predicted with any likelihood score or which are exchange reactions for dead-end metabolites) are each assigned by default a penalty of 1. The following MILP formulation then identifies a subset of reactions from the global set of candidate gap-filling reactions that together have the smallest sum of penalties and enable a minimum production of biomass (*β*=0.1 h^−1^ by default).

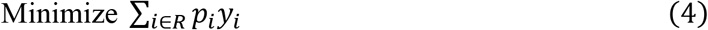

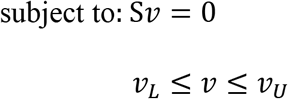

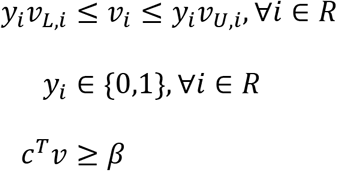

Here, the variables are the flux vector *v* and *y*. Each *y_i_* serves as an indicator variable: *y_i_* = 1 if and only if the *i*^th^ candidate gap-filler (with flux *v_i_*) is included in the solution.

### 2.4 Comparisons of Annotations and Networks

Architect was used to annotate enzymes to the proteomes of three species: *Caenorhabditis elegans*, *Escherichia coli* (strain K12) and *Neisseria meningitidis* (strain MC58) using sequences collated from WormBase (WS235, (Lee et al., 2018)), SwissProt and the Ensembl database (Howe et al., 2021) respectively. For each species, the naïve Bayes-based ensemble method was retrained by excluding sequences from the respective organism. Architect predictions were evaluated against gold standard datasets derived from UniProt for *C. elegans* and *N. meningitidis*, and SwissProt for *E. coli*. Performance was reported in terms of specificity and micro-averaged precision and recall (that is, irrespective of enzyme class size) (Sokolova & Lapalme, 2009).

For network comparisons, models generated by Architect, CarveMe and PRIAM (Claudel-Renard et al., 2003; Machado et al., 2018) were evaluated against two sets of gold-standard as described next. Performance was computed using micro-averaged precision and recall first using as a gold-standard, enzyme annotations assigned to genes in previously generated curated metabolic models: *C. elegans* – WormJam (Witting et al., 2018), *E. coli* – iML1515 (Monk et al., 2017) and *N. meningitidis* – Nmb_iTM560 (Mendum et al., 2011). As a second measure of performance, we compared the annotations included in the models following gap-filling against those in UniProt, here restricting the comparison to those ECs present in the relevant reaction database.

Finally, for *N. meningitidis* and *E. coli*, *in silico* knockout experiments were performed using the models generated by Architect and CarveMe. Genes predicted to be essential in these *in silico* experiments were subsequently compared to the results of two genome-scale knockout studies (Mendum et al., 2011; Monk et al., 2017). Since Architect, unlike CarveMe, does not predict complex gene-protein-reaction relationships (Thiele & Palsson, 2010), only those reactions associated with a single protein could be assessed through gene knockout experiments, where the flux through such reactions corresponding to a single protein was constrained to be zero.

## 3. RESULTS

### 3.1 Ensemble Methods Improve Enzyme Annotation

The motivation for developing an ensemble enzyme classifier is driven by the hypothesis that different enzymes (as defined by EC numbers) may be better predicted by different tools and, hence more accurate annotations may be obtained by combining predictions from individual tools. Based on this hypothesis we developed a novel enzyme prediction and metabolic reconstruction pipeline we term Architect (**Figure 1**). In brief, the pipeline begins with the prediction of enzyme annotations from proteome data (Module 1) using an ensemble classifier that combines predictions from five enzyme annotation tools (DETECT, EnzDP, Catfam, PRIAM and EFICAz). Next, these predictions of enzyme classification numbers are used to construct a functional metabolic model capable of generating biomass required for growth (Module 2).

**Figure 1:**
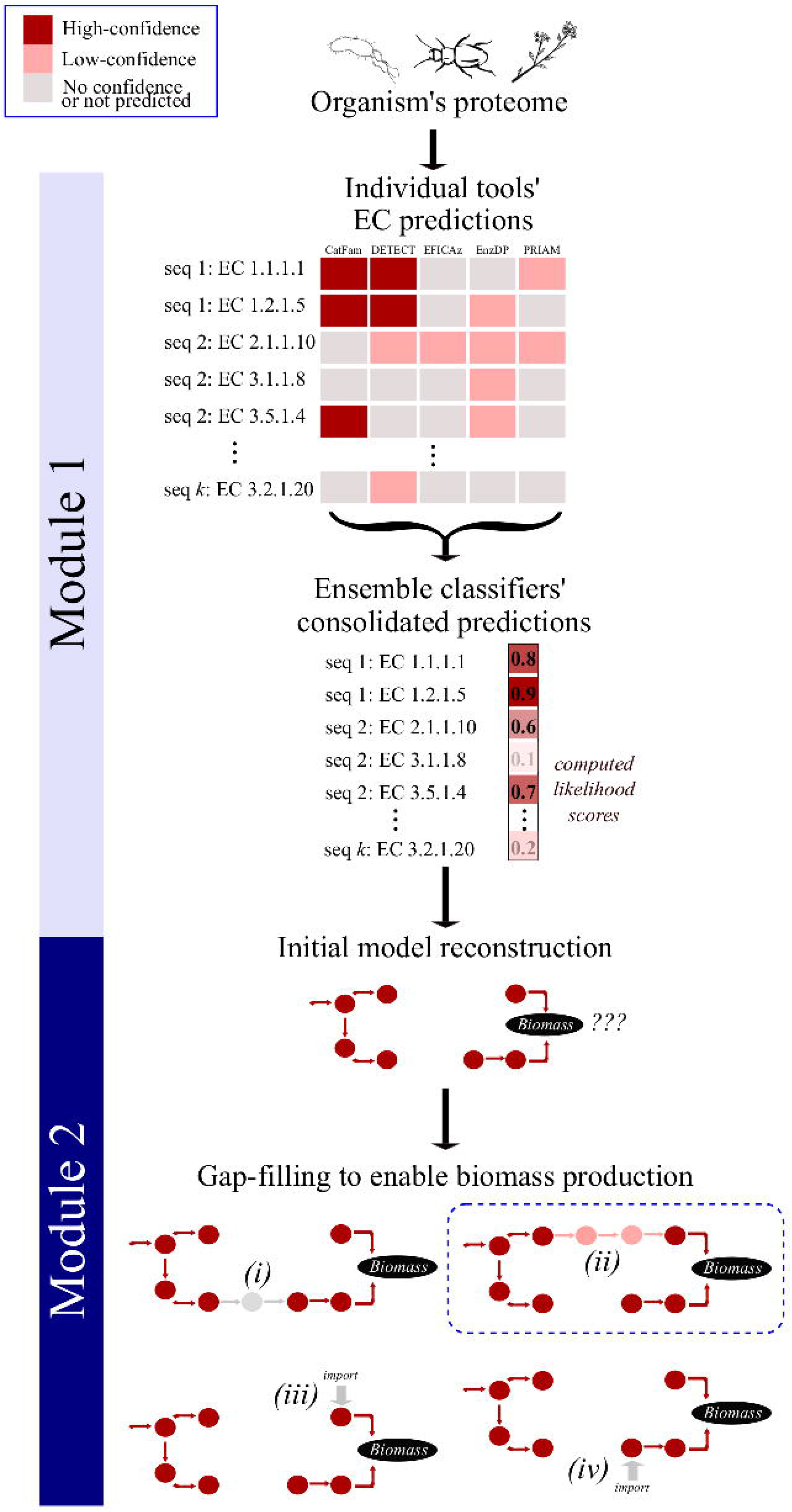
Overview of Architect’s methodology. Given an organism’s protein sequences, Architect first runs 5 enzyme annotation tools, then computes a likelihood score for each annotation using an ensemble approach (module 1). From high-confidence EC predictions, Architect reconstructs a high-confidence metabolic model which it then gap-fills to enable biomass production using the aforementioned confidence scores (module 2). In the illustrated example, 4 sets of reactions are considered for gap-filling, with the solution highlighted in the blue box yielding the highest score.

To test our hypothesis, we compared the performance of individual tools of interest and of various ensemble methods on a dataset of enzymatic sequences. Here we investigated three methods: naïve Bayes, logistic regression and random forest and employed five-fold cross-validation in which ensemble methods were trained on 80% of the enzymes annotated in SwissProt. Once trained, the performance of each classifier was tested on the remaining annotated enzymes by evaluating their high-confidence predictions against the database’s annotations. In addition to the classifiers and individual tools, we also examined the performance of a ‘majority rule’ approach (in which we assign an EC label to a protein on the basis of voting among the five tools), as well as an ‘EC-specific best tool’ approach (in which we assign an EC label to a protein based on best performing tools for that EC as seen in training). The performance of each dataset was computed using macro-averaged precision, recall and F1 to ensure that smaller EC classes were equally represented; performance on the non-enzymatic dataset (i.e. protein sequences not associated with either complete or partial EC annotations) is computed using specificity (see **Supplementary Material**).

Overall, we found that, with the exception of the majority rule, ensemble methods outperformed individual tools, resulting in both higher precision and recall (**Figure 2**). For example, the highest precision and recall of the individual tools—obtained by DETECT and PRIAM respectively—are lower than most of the ensemble methods applied. Indeed, except for majority rule, most ensemble methods perform similarly on the entire test set, as well as on subsets of test sequences with lower sequence similarity to training sequences (**Supplemental Figure 2**). Additionally, macro-recall on multifunctional proteins is decreased for the naïve Bayes, logistic regression and random forest classifiers when applying a heuristic which filters out predicted ECs other than the top-scoring EC and frequently co-occurring enzymes as seen in the training data (**Supplemental Figures 4 and 5** and **Supplemental Text**); therefore, henceforth, we evaluate performance of these classifiers by considering all their high-confidence EC predictions.

**Figure 2:**
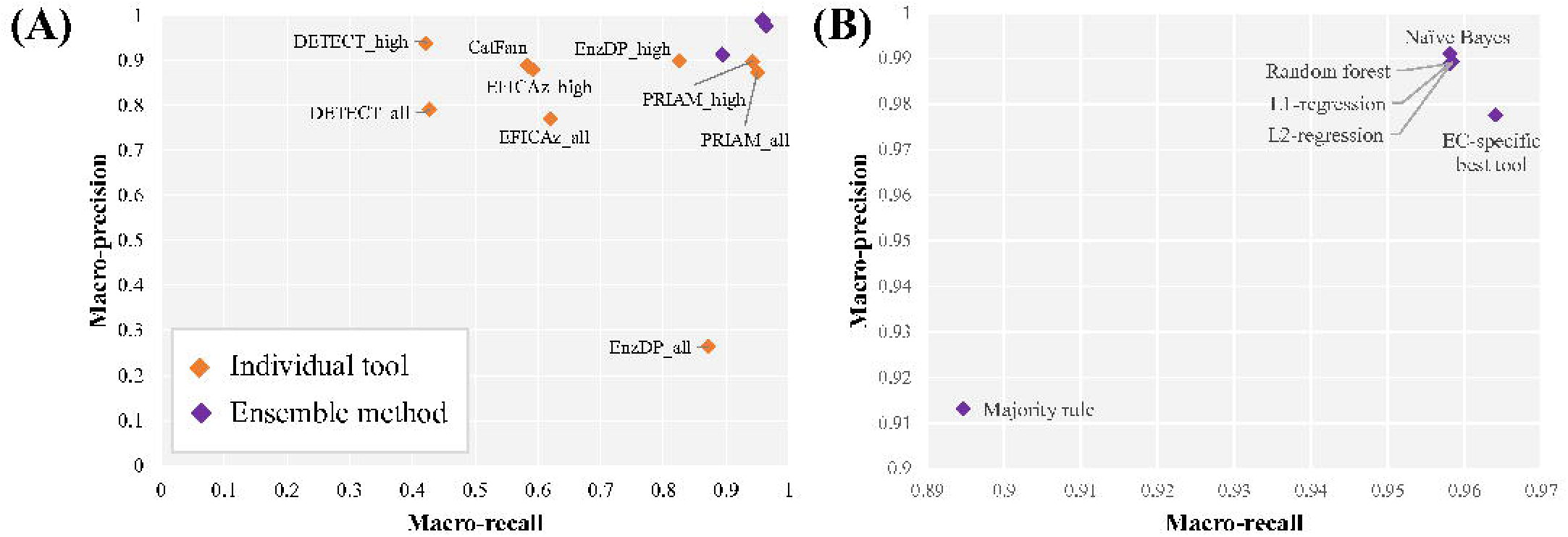
Performance of individual and ensemble enzyme annotation tools. (A) Precision/recall graph indicating performance of each method on the enzymatic test set from SwissProt, (B) focus on the improved performance of the ensemble methods. Our results show that combining predictions using almost any ensemble method gives better performance than using any individual tool.

Next, we consider the possibility that higher predictive range (defined as the number of ECs that a tool can predict) primarily drives the increased performance of the ensemble methods. Indeed, the ensemble approaches are superior when quantifying performance on ECs predictable by at least 2 tools (**Supplemental Figure 3**) but have similar precision and recall as DETECT on sequences annotated with ECs predictable by all tools (**Supplemental Figure 4**). However, when looking at DETECT’s or PRIAM’s class-by-class performance on ECs that they can predict, the ensemble method has higher precision and recall on more ECs than either DETECT (better precision on 176 versus 18 ECs and better recall on 116 versus 37 ECs) or PRIAM (better precision on 98 versus 38 ECs and better recall on 191 versus 41 ECs) (**Supplemental Figure 6**).

Given the main application of Architect is to annotate enzymes to an organism’s proteome, we were interested in assessing the ability of the ensemble approaches to minimize false positives. Applied to a set of proteins without EC annotations in SwissProt, we found that only the Naïve Bayes classifier gave comparable specificity as the individual tools (**Supplemental Figure 8**). Given the slightly elevated performance in terms of precision (for the enzymatic dataset) and specificity (for the non-enzymatic dataset), we chose the naïve Bayes classifier as the preferred method for Architect. We next investigated the performance of Architect to annotate the proteomes of three well-characterized organisms (*C. elegans*, *N. meningitidis* and *E. coli*; **Supplemental Figure 9**). We consider those annotations that feed into Architect’s model reconstruction module and compare them against high-confidence predictions by DETECT, EnzDP and PRIAM alone, these tools chosen due to their performance on the enzymatic dataset. For all three species, Architect yields both higher precision and recall than DETECT, and higher recall than EnzDP. In *C. elegans*, Architect gives higher recall than either PRIAM or EnzDP, albeit at the expense of precision and specificity. Overall, these results demonstrate Architect’s wider applicability to annotate specific organisms.

Finally, we investigated whether combining predictions from all 5 tools is required to obtain improved performance with respect to the individual tools. To this effect, we built the naïve Bayes-based method using predictions from fewer tools, then calculated performance once again on the held-out test set (**Supplemental Figure 7**). We observe that this procedure has a greater impact on macro-recall than macro-precision. In particular, leaving out predictions from both tools with the highest predictive ranges (EnzDP and PRIAM) had the greatest impact, while the F1-score decreased least when the tools with the lowest predictive ranges (CatFam and DETECT) are not included in the classifier. We also find that different tools are complementary to each other. While performance is mostly unaffected by excluding predictions from any single tool, combining predictions from at least 2 tools improves performance compared to using any single tool’s predictions. Indeed, simply combining predictions from PRIAM and any other tool yields better macro-precision than any tool in isolation. Intriguingly, training on predictions from EnzDP and PRIAM results in the highest performance among pairs of tools, with macro-precision showing only minimal increases with the inclusion of any other tool’s predictions. These results suggest that a user may obtain reasonably improved performance by combining predictions from fewer than 5 tools, for example, by excluding tools with longer running times (e.g. EFICAz (Ryu, Kim, & Lee, 2019)).

### 3.2 Automated Metabolic Reconstruction Using Architect

In addition to predicting suites of enzymes from an organism’s proteome, Architect utilizes these predictions to automatically reconstruct a functional metabolic model capable of generating biomass required for growth (see Methods). The process begins by querying the set of EC activities predicted by module 1, against a database of known reactions (either the Kyoto Encyclopedia of Genes and Genomes; KEGG (Kanehisa et al., 2021) or the Biochemical, Genetic and Genomic knowledgebase; BiGG (King et al., 2016), to construct an initial high confidence metabolic network. Next a gap filling algorithm is applied to identify enzymes absent from the model (potentially arising from uncharacterized enzymes or sequence diversity (Atkinson, Babbitt, & Sajid, 2009; Lespinet & Labedan, 2006)) to ensure pathway functionality and the ability of the model to generate biomass.

To validate the model reconstruction strategy, Architect was applied to reconstruct metabolic models for the three species previously investigated (*C. elegans*, *N. meningitidis* and *E. coli*) and compared to previously curated metabolic models (Mendum et al., 2011; Monk et al., 2017; Witting et al., 2018). For comparison, metabolic models were also generated using CarveMe (Machado et al., 2018) and PRIAM (Claudel-Renard et al., 2003). CarveMe performs metabolic model reconstruction in a top-down manner, retaining in a final functional model those reactions from the BiGG database (King et al., 2016) predicted with higher confidence scores for the organism of interest and required for model functionality; the confidence score is by default computed based on sequence similarity (Buchfink et al., 2015). On the other hand, PRIAM uses its high-confidence predictions to output a KEGG-based metabolic model. To account for differences that may be introduced from using different databases of reactions, two versions of Architect models were predicted using either the KEGG or the BiGG database.

Focusing on the KEGG-based reconstructions (**Figure 3 and Supplemental Figure 10**), Architect produced models of higher precision than either PRIAM or CarveMe for *C. elegans* and *N. meningitidis*, and higher recall for *C. elegans* alone. However, in *E. coli*, CarveMe has significantly better precision and recall than Architect, likely a result of the presence of curated *E. coli*-related reactions in the BiGG database used for its reconstructions. Similarly, Nmb_iTM560 was based on the *i*AF1260 *E. coli* model (Feist et al., 2007), again likely contributing to CarveMe’s higher recall when reconstructing a *N. meningitidis* metabolic model. Indeed, when the models are compared against organism-specific datasets retrieved from UniProt, Architect has higher precision and recall than either CarveMe or PRIAM for *E. coli* (as well as higher precision and recall than CarveMe for the other two species). This highlights differences in annotations associated with the curated models and UniProt. We also note that higher EC coverage in KEGG compared to BiGG (**Supplemental Figure 11**) may contribute towards higher recall by Architect and PRIAM compared to CarveMe, a factor we account for by next using the BiGG database for Architect’s model reconstructions.

**Figure 3:**
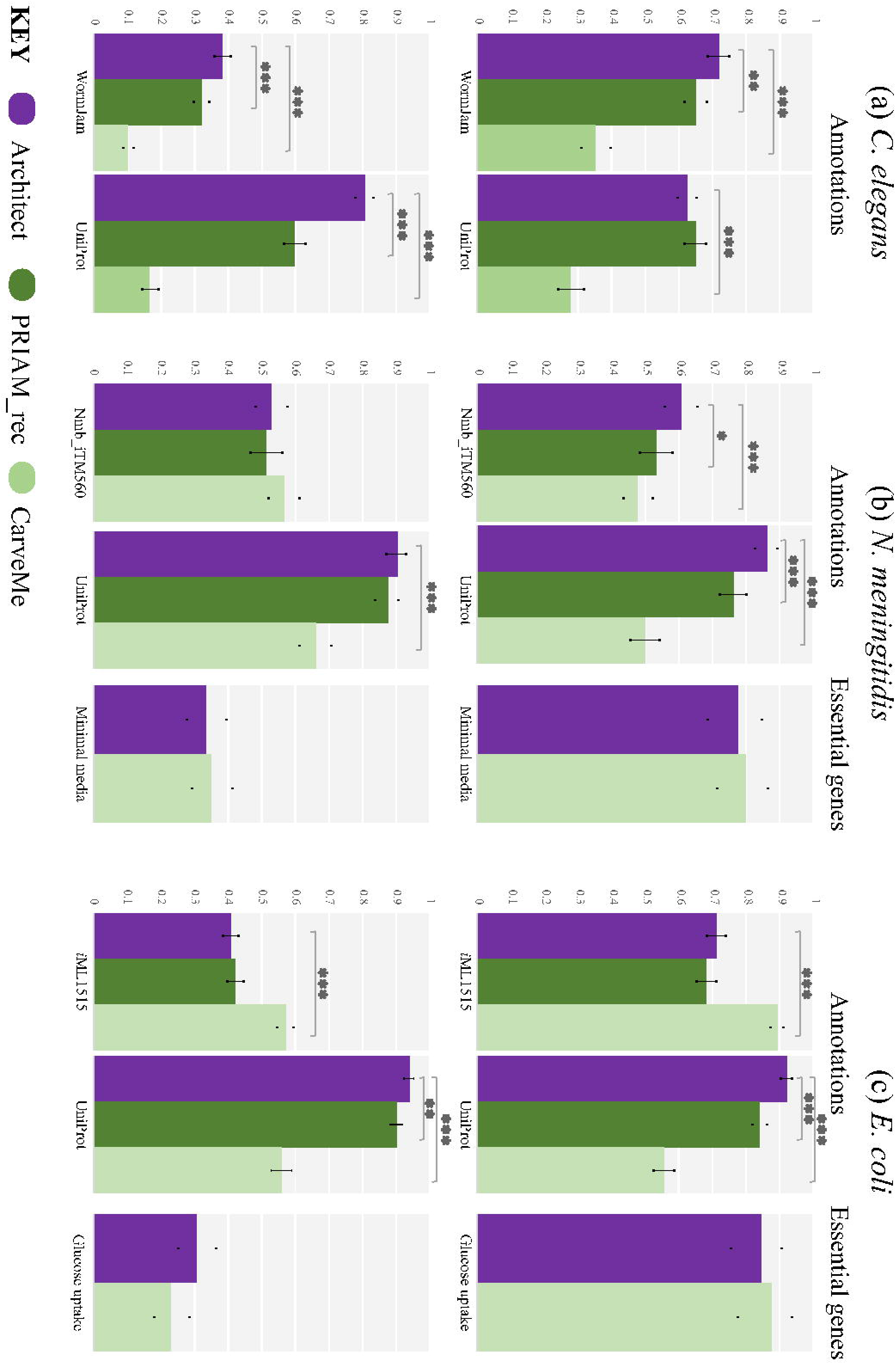
Performance of Architect as a model reconstruction tool versus CarveMe and PRIAM (as a model reconstruction tool). Quality of annotations is computed against the curated models over the genes found in these models, and against UniProt/SwissProt when restricting to those sequences found in the database and with ECs present in the KEGG reaction database. Genes determined essential *in silico* are compared against those whose knockout was tested *in vivo*. Error bars show the 95% confidence interval for precision and recall, each considered as the estimate of a binomial parameter. P-values, computed using Fisher’s exact test, are calculated only between Architect and either CarveMe or PRIAM (with *, ** and *** representing *p* less than 0.05, 0.005 and 0.0005 respectively).

Turning to models constructed with the BiGG database, as for the KEGG-based models, we find that Architect has higher precision for both *C. elegans* and *N. meningitidis*, and higher recall for the former (**Supplemental Figure 12**). However, we now observe similar precision in *E. coli*, consistent with the reliance on the BiGG database, which avoids the inclusions of ECs exclusive to the KEGG database (and hence absent in *i*ML1515). As expected from the lower coverage of ECs provided by the BiGG database, recall differences in *E. coli* qualitatively remain unchanged while in *N. meningitidis*, Architect’s recall, sacrificed for higher precision, is now significantly lower. Again, to account for the construction of the BiGG database from previously curated metabolic reconstructions that include *E. coli*, we compared the protein-EC annotations in the reconstructed models to those found in UniProt. Architect is then perceived as having higher precision in both *N. meningitidis* and *E. coli*, and greater recall in *E. coli*. Thus, our results indicate that Architect may be used to produce models with more accurate EC annotations than either CarveMe or the PRIAM-reconstruction tool. Additionally, the choice of reaction database when running Architect may impact the range of ECs covered. Indeed, it should be noted that the BiGG reaction database used here contains reactions from bacterial models only; thus, use of KEGG for eukaryotic reconstructions is more appropriate, as evidenced by the better recall with the *C. elegans* metabolic model.

### 3.3 Metabolic Reconstructions Benefit from Annotation Tools with High Predictive Range

From the previous comparisons of metabolic reconstructions, it is clear that there is a difference between Architect’s performance as an enzyme annotation tool and as a tool for model reconstruction. This raises the question of whether improvement in enzyme annotation is associated with a corresponding improvement in accuracy of model reconstruction. To address this, Architect’s model reconstruction module was applied to high-confidence predictions from individual tools, rather than from the naïve Bayes-based method (**Supplemental Figure 13**). Overall, models constructed from high-confidence DETECT predictions resulted in significantly lower recall compared to either the curated models or UniProt annotations. This is consistent with the idea that high predictive range is an important attribute in an enzyme annotation tool used for model reconstruction; accordingly, when comparing against model annotations, using high-confidence predictions from either EnzDP or PRIAM instead of DETECT resulted in models with higher recall. At the same time, the resulting recall does not significantly differ from the recall obtained when using predictions from the ensemble method. Furthermore, similar precision is obtained when using EnzDP, PRIAM or the ensemble method when reconstructing *C. elegans* or *N. meningitidis* models; in the case of *E. coli*, significantly better precision is obtained by using predictions from EnzDP instead of the ensemble method. Therefore, based on comparisons of EC annotations derived from curated models, there appears to be no benefit to substituting predictions from tools with high predictive range with those from the ensemble approach. However, when comparing against annotations obtained from UniProt, the ensemble method results in higher recall than EnzDP for all 3 organisms and higher recall than PRIAM for *C. elegans*; similar precision is observed in all organisms, with the exception of lower precision than EnzDP and PRIAM for *C. elegans*. These conflicting results reflect inherent differences in the gold-standard datasets, which in turn may indicate that existing curated models may have potential to be further expanded by using UniProt annotations. Interestingly, use of PRIAM’s high confidence EC predictions as input to Architect’s reconstruction module results in more accurate annotations than models generated from PRIAM’s reconstruction tool (**Figure 3 and Supplemental Figure 13**), highlighting methodological differences between the two tools. It should also be noted that unlike the PRIAM pipeline, models constructed by Architect are simulation-ready.

## 4. CONCLUSION AND FUTURE DIRECTIONS

Here, we present Architect, an approach for automatic metabolic model reconstruction. The tool consists of two modules: first, enzyme predictions from multiple tools are combined through a user-specified ensemble approach, yielding likelihood scores which are then leveraged to produce a simulation-ready metabolic model. Through the use of various gold-standard datasets, we have shown that Architect’s first module produces more accurate enzyme annotations, and that its second module can be used to produce organism-specific metabolic models with better annotations than similar state-of-the-art reconstruction tools, including CarveMe and PRIAM. Our expectation is that these models serve as near-final drafts, requiring users to perform only minimal curation to incorporate organism-specific data. For example, models for eukaryotic organisms may require the independent definition of cellular compartments. Interestingly, it is unclear whether improvements in enzyme annotation, other than in terms of predictive range, lead to the construction of models with either improved annotations or greater accuracy of simulations. Instead, we propose three improvements to the input and the algorithm of the model reconstruction module that will likely yield better models. First, we find that most essential genes also incorporated into the final models were not predicted to be essential *in silico* (see **Supplemental Text**), suggesting that more accurate predictions of gene essentiality may be obtained by better encoding gene-protein-reaction relationships or by limiting the reactions included in the high-confidence model based on EC annotation. Second, improving the predictions of transport reactions is needed to define the accurate import and export of metabolites which otherwise represent dead-ends in the initial network; in turn, this may lead to fewer blocked reactions (given high proportions of inactive reactions as shown in **Supplemental Table 2**). Third, considerations of thermodynamics have been absent from our reconstruction pipeline, whether in terms of reaction reversibility, or in terms of gap-filling. Identifying thermodynamically likely solutions for gap-filling is expected to result in more biologically realistic models (Fleming, Thiele, & Nasheuer, 2009).

## Acknowledgements

We thank members of the Parkinson and Moses labs for constructive feedback throughout the course of this project.

## SUPPLEMENTAL FIGURES

**<u>Supplemental Figure 1: Properties of ECs and proteins included in Architect’s database</u>**

(A): Proportion of ECs in Architect’s training and test datasets combined associated with different numbers of proteins, with proportions relevant to the subset of those ECs predictable by all tools indicated in grey. The percentages of ECs predictable by any tool and associated with different numbers of proteins are indicated above the bars.

(B): Intersection of ECs predictable by different tools, when focussing on ECs (i) found in Architect’s training database and (ii) more generally. These Venn diagrams were made using the interface at http://bioinformatics.psb.ugent.be/webtools/Venn.

(C): Number of proteins in the training and test sets combined associated with 1 or more ECs. The percentage of the proteins involved in each category is given above the bars.

**<u>Supplemental Figure 2: Performance of different ensemble methods on test set at different levels of sequence identity to the training data</u>**

Comparison of macro-averaged (A) precision, (B) recall, and (C) F1-score of ensemble methods at different MTTSIs—defined in the supplemental text.

**<u>Supplemental Figure 3: Performance of different ensemble methods on test set when considering ECs predictable by an increasing number of tools</u>**

Comparison of (A) macro-precision, (B) macro-recall, and (C) F1-score of individual tools and ensemble methods on proteins with ECs predictable by at least 1, 2, 3, 4 tools and by all tools.

**<u>Supplemental Figure 4: Precision and recall of ensemble methods and individual tools for enzyme annotation with respect to ECs predictable by all tools and on different sets of proteins</u>**

(i) Comparison of precision and recall of ensemble (with and without multi-EC filtering) and individual tools on ECs predictable by all tools in (A) the entire test data set, (B) only proteins with a single EC, and (C) only multifunctional proteins. The subscripts “all” and “high” reflect whether all or only high-confidence predictions from individual tools were considered, respectively.

(ii): Comparison of precision and recall of ensemble methods (with and without multi-EC filtering) on ECs predictable by all tools in (A) the entire test data set, (B) only proteins with a single EC, and (C) only multifunctional proteins.

**<u>Supplemental Figure 5: Precision and recall of ensemble methods and individual tools for enzyme annotation with respect to ECs predictable by any tool and on different sets of proteins</u>**

(i) Comparison of precision and recall of ensemble (with and without multi-EC filtering) and individual tools on ECs predictable by any tool in (A) the entire test data set, (B) only proteins with a single EC, and (C) only multifunctional proteins. The subscripts “all” and “high” reflect whether all or only high-confidence predictions from individual tools were considered, respectively.

(ii): Comparison of precision and recall of ensemble methods (with and without multi-EC filtering) on ECs predictable by any tool in (A) the entire test data set, (B) only proteins with a single EC, and (C) only multifunctional proteins.

**<u>Supplemental Figure 6: Class-by-class comparison of performance of the naïve Bayes method and two enzyme annotation tools, DETECT and PRIAM</u>**

(A): Class-by-class comparison of (i) precision and (ii) recall between the naïve Bayes-based ensemble method and DETECT (high-confidence) when considering ECs predictable by DETECT. Each dot represents an EC class, and those dots above the line correspond to EC classes for which the ensemble method performs better. Only EC classes with defined precision from both DETECT and the ensemble classifier are shown.

(B): Same as (A), except that the class-by-class comparison is between the naïve Bayes-based ensemble method and PRIAM (high-confidence predictions) and on ECs predictable by PRIAM. Again, only EC classes with defined precision from both PRIAM and the ensemble classifier are shown.

**<u>Supplemental Figure 7: Comparison of performance of the naïve Bayes method on the test set when training on predictions from combinations of fewer than 5 tools.</u>**

In (A), the light green square in each row indicates when a tool’s predictions are being considered in a combination; this table shows the combinations of tools ranked from highest F1-score to lowest amongst those involving 2, 3 and 4 tools respectively. In (B), the purple diamond indicates the performance when the naïve Bayes-based method is trained on the entire set of predictions, whereas each green circle indicates performance when considering a particular combination of tools (number corresponding to the rank in (A)).

**<u>Supplemental Figure 8: Specificity of individual tools (high-confidence) and ensemble approaches on non-enzymatic dataset</u>**

**<u>Supplemental Figure 9: Comparison of organism-specific performance of Architect’s enzyme annotation tool using predictions by the naïve Bayes ensemble method and three individual tools, DETECT, EnzDP and PRIAM</u>**

High-confidence PRIAM predictions were included to the predictions of the naïve Bayes classifiers for those ECs outside of Architect’s predictive range. The comparison was done over those proteins present in both the input protein sequence file and those found in UniProt/SwissProt. Error bars show the 95% confidence interval for precision, recall and specificity, each considered as the estimate of a binomial parameter. P-values are calculated using Fisher’s exact test and between Architect results and other tools only (with *, ** and *** representing *p* less than 0.05, 0.005 and 0.0005 respectively).

**<u>Supplemental Figure 10: Overlap of annotations in models reconstructing with Architect (based on KEGG), CarveMe and PRIAM’s reconstruction tool using as gold-standard (i) curated models, and (ii) UniProt/SwissProt.</u>**

**<u>Supplemental Figure 11: Number of pathway-specific ECs present in Architect’s KEGG database versus in CarveMe’s main BiGG database.</u>**

Each dot represents a pathway in KEGG, and the diagonal line indicates the points on the graph where a pathway is covered by the same number of ECs through KEGG and BiGG.

**<u>Supplemental Figure 12: Performance of Architect as a model reconstruction tool (using BiGG as the reaction database) versus CarveMe in the case of the three organisms of interest.</u>**

Quality of annotations is computed against the curated models over the genes found in these models, and against UniProt/SwissProt when restricting to those sequences found in the database and with ECs present in the BiGG reaction database. Genes determined essential *in silico* are compared against those whose knockout was tested *in vivo*. Error bars show the 95% confidence interval for precision and recall, each considered as the estimate of a binomial parameter. P-values are calculated using Fisher’s exact test (with *, ** and *** representing p less than 0.05, 0.005 and 0.0005 respectively).

**<u>Supplemental Figure 13: Performance of Architect as a model reconstruction tool when using as input high-confidence predictions from the naïve Bayes-based method or from DETECT, EnzDP or PRIAM.</u>**

Quality of annotations is computed against the curated models over the genes found in these models, and against UniProt/SwissProt when restricting to those sequences found in the database and with ECs present in the BiGG reaction database. Genes determined essential *in silico* are compared against those whose knockout was tested *in vivo*. Error bars show the 95% confidence interval for precision and recall, each considered as the estimate of a binomial parameter. P-values, computed using Fisher’s exact test, are calculated only between Architect run on the ensemble method and on Architect run on any of the individual tools’ predictions (with *, ** and *** representing *p* less than 0.05, 0.005 and 0.0005 respectively).

