## Supplemental Text for "Architect: a tool for producing high-quality metabolic models through improved enzyme annotation"

### Supplementary information

### Table of contents


| Page | Section |
|---|---|
| 2 | **A. Individual enzyme annotation tools** |
| 2 | **B. Ensemble approaches** |
| 2 | Majority rule |
| 3 | EC-specific tool |
| 4 | Naïve Bayes |
| 6 | Training on different tool combinations |
| 6 | Logistic regression model |
| 7 | Random forest |
| 8 | **C. Methods of analyzing results** |
| 8 | Performance measures |
| 9 | Performance on multi-functional proteins |
| 9 | Performance on test set based on sequence similarity to training data |
| 10 | **D. Metabolic model reconstruction** |
| 10 | KEGG database: curation |
| 11 | BiGG databases and addition of non-EC associated reactions via sequence similarity |
| 11 | Relevant simulation details |
| 12 | Identification of gap-filling candidates required for biomass production |
| 13 | Essentiality experiments |
| 13 | **E. Model reconstruction: technical details** |
| 15 | **F. Additional results** |
| 15 | 3.1a Filtering multifunctional enzyme predictions has minimal impact on performance |
| 15 | 3.4 Comparing gene essentiality results |
| 16 | **G. Supplementary Tables** |
| 16 | 1: Overlap of *in silico* determined essential genes with those found essential *in vivo*. |
| 17 | 2: Comparisons of various aspects of model reconstruction for *C. elegans*, *N. meningitidis* and *E. coli*. |
| 19 | 3: Breakdown of annotations of SwissProt sequences by individual and ensemble methods into true positives, true negatives, false positives and false negatives |
| 20 | 4: Breakdown of organism-specific annotations by ensemble and individual tool into true positive, false positive and false negative. |
| 21 | **H. Bibliography** |

### A. Individual enzyme annotation tools

We ran the following enzyme annotation tools: EFICAz v2.5.1 [1], PRIAM [2], DETECT v2 [3], EnzDP [4] and CatFam [5]. For each tool, we only considered complete EC annotations (that is, of the form x.x.x.x, where x is a number). We labelled as "high-confidence" predictions that satisfy the following criteria for each tool: those predictions labelled as "high-confidence" in EFICAz; those achieving a score above the cutoff of 0.2 and 0.7 in PRIAM and EnzDP respectively; and those passing the EC-specific cutoffs of DETECT v2. The cutoff of 0.7 was found optimal when looking at EnzDP's performance on the training data (**Figure 1** below), while the 0.2 cutoff was suggested in the README file associated with PRIAM for sequences of non-bacterial origin. We considered all predictions from CatFam to be of high-confidence. Remaining predictions from EFICAz were considered to be of "low-confidence". For the other 3 tools, remaining predictions with a score of at least 0.0001 were considered "low-confidence" predictions. These levels of confidence are used in feature vector construction when training and testing the ensemble methods. In general, each element of a feature vector represents the level of confidence in a tool's prediction.

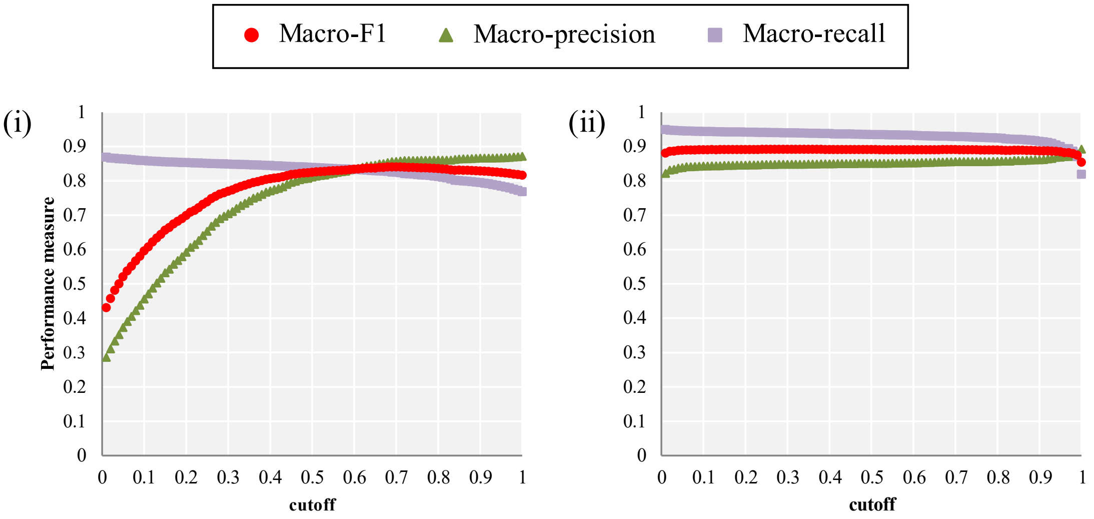


*Figure 1*: Macro-precision, macro-recall and F1-score of (i) EnzDP and (ii) PRIAM at different cutoffs. Performance is summarized over 80% of Architect's entire training database; this corresponds to the portion of the database used to train the ensemble methods which were then tested with performance summarized in **Figure 1** of the main text.

### B. Ensemble approaches

#### *Majority rule*

One of the simplest benchmarking methods we employ to combine predictions from multiple classifiers is through a majority rule. Here, we explore three main voting schemes: (M1) simple majority, (M2) plurality voting [6], and (M3) unanimous voting. When a simple majority rule (M1) is applied, an EC is assigned to a protein if it is predicted by at least half (that is 3) of all tools applied. In plurality voting (M2), a protein is assigned the EC predicted by most methods; this is a more relaxed version of M1 such that agreement by fewer than 3 tools for an EC is allowed. On the other hand, unanimous voting (M3) is the most conservative of the majority rules, where an EC is assigned to a protein sequence only if all 5 enzyme annotation tools agree. In another version of the M1 and M3 rules (M1.5, M3.5), rather than considering 5 as the "maximum" number of tools, we define a different "maximum" for each EC, counting it as the total number of tools that can actually predict that EC. Furthermore, we present results of experimenting with only high-confidence predictions from each method, or on all predictions. We note that both in cross-validation and on the test data (Table 1), applying M2 on the high-confidence dataset gives the best result, and therefore present results from this rule unless otherwise specified.

Table 1: Performance of various voting rules for enzyme annotation in cross-validation and on test set.
The suffixes "_low" and "_high" indicate whether both all or only high-confidence predictions were considered respectively.

| Flavour of voting rule | Averaged macro-F1 over cross-validation | Macro-F1 on test set |
|---|---|---|
| M1_low | 82.5% | 82.0% |
| M1.5_low | 67.6% | 67.3% |
| M2_low | 84.0% | 80.9% |
| M3_low | 41.8% | 41.8% |
| M3.5_low | 78.7% | 77.9% |
| M1_high | 82.1% | 81.7% |
| M1.5_high | 90.4% | 89.5% |
| M2_high | 90.7% | 90.4% |
| M3_high | 39.9% | 39.8% |
| M3.5_high | 77.6% | 77.0% |

#### *EC-specific tool*

Another benchmarking method is to find, for each EC, the tool(s) which performs best in the training set, and to retain predictions of that EC in the test set only when made by the best-performing tool(s). We define the best-performing tool as the tool that achieves the highest F1-score on the training data. To control for differences across the data from influencing what defines the top-performer for an EC, we find the tool(s) achieving the highest F1-measure over each four-fifths of the training data in cross-validation. A method that is the top-performer for an EC across all 5 four-fifths of the data is marked as a "high-confidence EC-specific tool", and is otherwise marked as a "low-confidence EC-specific tool" (given that it is the top performer for some but not all parts of the training data).

We specifically considered the confidence with which each tool makes a prediction to identify EC-specific best tools. Therefore, for an EC $x$, we construct feature vectors ($v$) to indicate whether an annotation tool predicts EC $x$ (whether with low- or high-confidence), and further indicate (with an additional feature) if the prediction is made with high-confidence. For each prediction of EC $x$:

$$v_1 = \begin{cases} 1 \text{ if CatFam has made this prediction} \\ 0 \text{ otherwise} \end{cases} \quad (1)$$

$$v_2 = \begin{cases} 2 \text{ if DETECT has made this prediction with high-confidence} \\ 1 \text{ if DETECT has made this prediction} \\ 0 \text{ otherwise} \end{cases}$$

$$v_3 = \begin{cases} 2 \text{ if EFICAz has made this prediction with high-confidence} \\ 1 \text{ if EFICAz has made this prediction} \\ 0 \text{ otherwise} \end{cases}$$

$$v_4 = \begin{cases} 2 \text{ if EnzDP has made this prediction with high-confidence} \\ 1 \text{ if EnzDP has made this prediction} \\ 0 \text{ otherwise} \end{cases}$$

$$v_5 = \begin{cases} 2 \text{ if PRIAM has made this prediction with high-confidence} \\ 1 \text{ if PRIAM has made this prediction} \\ 0 \text{ otherwise} \end{cases}$$

Thus, this algorithm can identify tools that perform best for an EC only when their high-confidence predictions are considered.

Using predictions from "low-confidence EC-specific tools" (that is, top performers on at least a subset of the data) gives the best results on the test data, and therefore we present results from this technique unless otherwise specified (Table 2).

Table 2: Performance of the different flavours of the EC-specific best tool on test set

| Applying EC-specific best tools that are found: | Macro-F1 on test set |
|---|---|
| on at least a subset of training data | 97.1% |
| on all subsets of training data | 94.8% |

***Naïve Bayes***

We built a naïve Bayes classifier for each EC class. For a particular EC predicted for a protein, we calculate the likelihood score in the prediction as follows, where $n$ here is the number of tools that can predict the EC:

$$p(y = 1 | v_1 = \alpha_1, v_2 = \alpha_2, \ldots, v_n = \alpha_n) = \frac{p(y = 1) \cdot \prod_{i=1}^{n} p(v_i = \alpha_i | y = 1)}{\sum_{k \in \{0,1\}} p(y = k) \cdot \prod_{i=1}^{n} p(v_i = \alpha_i | y = k)} \quad (2)$$

$y$ is a binary variable denoting the membership of the protein in the EC class (true if and only if $y = 1$). We calculate $p(y = k)$ as follows:

$$p(y = 1) = \frac{\text{number of proteins actually in EC class}}{\text{number of proteins either predicted or actually in EC class}} \quad (3)$$

$$p(y = 0) = \frac{\text{number of proteins not actually in EC class (i.e., false positives)}}{\text{number of proteins either predicted or actually in EC class}} \quad (4)$$

$v_i$ represents the value of the feature vector of the $i^{th}$ tool. As per the nature of naïve Bayes, we make the assumption that the predictions of individual tools are conditionally independent from each other given the class label $y$. As detailed below, we experimented with different feature vector constructions corresponding to different methods of calculating $p(v_i | y = k)$.

**Approximating distribution of features using a Bernouilli distribution**

**(i) Considering high-confidence predictions only**

We only consider high-confidence predictions made by individual tools, such that the feature vectors are constructed as follows:

$$v_i = \begin{cases} 1 \text{ if the ith tool predicted this EC with high-confidence} \\ 0 \text{ otherwise} \end{cases} \quad (5)$$

*i* ranges from 1 to *m* where *m* is the number of tools that can predict the EC in question. Then:

$$p(v_i = \alpha_i | y = k) = \frac{(\text{number of proteins for which } v_i{=}\alpha_i \text{ and } y{=}k) + 1}{(\text{number of proteins for which } y{=}k) + 2} \quad (6)$$

where the constants added to the numerator and denominator are due to Laplace smoothing.

**(ii) Considering all predictions**

We consider all predictions made by individual tools, whether with low- or high-confidence:

$$v_i = \begin{cases} 1 \text{ if the ith tool predicted this EC} \\ 0 \text{ otherwise} \end{cases} \quad (7)$$

We use equation (6) as given above.

**(iii) Accounting for the level of confidence in tools' predictions**

We consider all predictions made by individual tools, and have each feature reflect the level of confidence with which the prediction is made by the corresponding tool. All CatFam predictions are considered of high-confidence. Therefore, for ECs predictable by CatFam:

$$v_1 = \begin{cases} 1 \text{ if CatFam predicted this EC} \\ 0 \text{ otherwise} \end{cases} \quad (8)$$

**For other tools that can predict the EC in question, we use the following:**

$$v_i = \begin{cases} 2 \text{ if the ith tool has made this prediction with high-confidence} \\ 1 \text{ if the ith tool has made this prediction with low-confidence} \\ 0 \text{ otherwise} \end{cases} \quad (9)$$

**Approximating distribution of continuous scores with a normal distribution, and categorical features with a Bernouilli distribution**

Here, for predictions made by DETECT, EnzDP and PRIAM, we assume that the actual confidence scores with which an EC is predicted follows a normal distribution. For each of these tools, we compute the sample mean ($\mu$) and the sample standard deviation ($\sigma$) of the scores for the corresponding proteins. Then, for a score *s*:

$$p(v_i = s | y = k) = \frac{1}{\sigma\sqrt{2\pi}} e^{-\frac{1}{2}\left(\frac{s-\mu}{\sigma}\right)^2} \quad (10)$$

CatFam and EFICAz predictions are not given with scores. Therefore, when assigning the value of the feature corresponding to these tools, we use equations (8) and (9) respectively.

We found, as part of cross-validation and testing, that while there was little difference between the performance of the different flavours of naïve Bayes (Table 3), using high-confidence predictions only (i.e. (i)) gives the highest performance. Therefore, results correspond to these classifiers are presented in the paper.

Table 3: Performance of various flavours of naïve Bayes classifiers in cross-validation and on test set. The results of applying the different flavours of naïve Bayes are indicated by the suffixes. ((i): "_high", (ii): "_all", (iii): "_categorical", and (iv): "_mixed")

| Flavour of naïve Bayes | Averaged macro-F1 over cross-validation | Macro-F1 on test set |
|---|---|---|
| bernouilli_all | 96.2% | 96.4% |
| bernouilli_high | 97.2% | 97.4% |
| bernouilli_categorical | 97.0% | 97.2% |
| bernouilli_mixed | 96.7% | 97.2% |

We would like to point out that in the case of the naïve Bayes classifier, as with the following algorithms, in cases where proteins with an EC *x* are found to never be falsely predicted in training, the algorithm assigns EC *x* whenever a prediction is made by the same tools found to never make a false positive prediction in training.

#### *Training on different tool combinations*

We experimented with training on predictions from different combinations (or subsets) of tools. Here we built Naïve Bayes classifiers using only high-confidence predictions from individual tools. The feature vectors for each EC are constructed as described in (6) with the following exceptions:

1. Predictions coming only from tools of interest are considered.
2. The length of the feature vector is now the number of tools in the combination of interest that can also predict the EC.

#### *Logistic regression model*

We train a logistic regression model for each EC class to examine the impact of using a weighted combination of classifiers. In this case, our feature vectors are constructed using one-hot encoding, reflecting—except in the case of CatFam—whether the EC was predicted and if so whether it was done with high- or low-confidence; in the case of CatFam, the encoding only reflects whether a prediction was made or not. As with naïve Bayes, we build the feature vector to be of a length reflecting the tools that can predict the EC in question.

We performed five-fold cross-validation to find the optimal value for parameter *C* when using either L1- or L2-regularization. The cost function for the logistic regression model we train for each EC is as follows in the case of L2-regularization

$$\frac{1}{2} w \cdot w + C \sum_{i=1}^{n} \log\left(e^{-y_i(X_i^T w + c)} + 1\right) \qquad (11)$$

and in the case of L1-regularization

$$\|w\|_1 + C \sum_{i=1}^{n} \log\left(e^{-y_i(X_i^T w + c)} + 1\right) \qquad (12)$$

Here, *n* is the number of training examples and *i* indexes over each of these examples. *w* is a vector for which $w_j$ is the weight for the $j^{\text{th}}$ feature; $X_i$ is the feature vector representing the $i^{\text{th}}$ training example; $y_i$ is the label of the $i^{\text{th}}$ training example where $y_i = 1$ if the $i^{\text{th}}$ protein belongs to the EC class and $y_i = 0$ otherwise. *C* represents the inverse of the regularization strength (that is, smaller values of $C$ result in higher regularization as the norm of the weight vector grows more important in the cost function).

We also compute regression models by applying a correction for class imbalance. Class imbalance arises given that for each EC, the number of positive examples (true positives and false negatives) is not typically comparable to the number of negative examples (EC falsely predicted by any tool). To do so, we used the "balanced" option in the SciKit-learn package [7, 8], which calculates class weights $\sigma_i$ for class $i$ as follows:

$$\sigma_i = \frac{\sum_{i=1}^{k} N_i}{k \cdot N_i} \quad (13)$$

Here, $k$ is the total number of classes ($k$=2 in this report), and $N_i$ is the number of training examples with label $i$. The class weight is applied to the cost function as follows in the case of L2-regularization

$$\frac{1}{2} w \cdot w + C \sum_{i=1}^{n} \sigma_i \cdot \log\left(e^{-y_i(X_i^T w + c)} + 1\right) \quad (14)$$

and in the case of L1-regularization

$$\|w\|_1 + C \sum_{i=1}^{n} \sigma_i \cdot \log\left(e^{-y_i(X_i^T w + c)} + 1\right) \quad (15)$$

As a consequence, the error in predicting the smaller class (either positive or negative examples for an EC) is amplified when minimizing the value of the cost function.

During cross-validation, we find the following values of C to give the highest macro-averaged F1-score. Therefore, we use the corresponding values on the test set.

Table 4: Optimal C-value as found through cross-validation for various settings of logistic regression.

| **Regularization** | **Balanced?** | **C** |
|---|---|---|
| L1 | No | 400 |
| L1 | Yes | 90 |
| L2 | No | 1000 |
| L2 | Yes | 10 |

#### *Random forest*

We train a random forest classifier [9] for each EC class. Each feature vector here is of length 5, each cell representing the confidence of the EC prediction by the corresponding tool.

$$v_1 = \begin{cases} 1 \text{ if CatFam has made this prediction} \\ 0 \text{ otherwise} \end{cases} \quad (16)$$

$$v_2 = \begin{cases} 2 \text{ if DETECT has made this prediction with high-confidence} \\ 1 \text{ if DETECT has made this prediction with low-confidence} \\ 0 \text{ otherwise} \end{cases}$$

$$v_3 = \begin{cases} 2 \text{ if EFICAz has made this prediction with high-confidence} \\ 1 \text{ if EFICAz has made this prediction with low-confidence} \\ 0 \text{ otherwise} \end{cases}$$

$$v_4 = \begin{cases} 2 \text{ if EnzDP has made this prediction with high-confidence} \\ 1 \text{ if EnzDP has made this prediction with low-confidence} \\ 0 \text{ otherwise} \end{cases}$$

$$v_5 = \begin{cases} 2 \text{ if PRIAM has made this prediction with high-confidence} \\ 1 \text{ if PRIAM has made this prediction with low-confidence} \\ 0 \text{ otherwise} \end{cases}$$

We built random forest classifiers optimized per EC as follows. We performed cross-validation for each EC, varying the seed to 3 values to account for the inherent randomness in the construction of random forest classifiers, and found the simplest settings of the hyper-parameters that provided the best F1-score. The hyper-parameters, sorted by decreasing priority, are as follows (with values indicated in parentheses): number of trees (10, 20, 30, 40, 50, 100), maximum tree depth (2, 4, 6, 8, 10), number of features to consider when performing a split (2 or 5), and whether the gini or the entropy function is considered to measure the quality of a split.

In each case, we also separately experimented with using two types of balanced random forest classifiers and a non-balanced random forest classifier. With the balanced random forest classifier (referred to simply as "balanced"), class imbalance is corrected when building the trees; in another flavor of the balanced classifier, the bootstrap sample upon which each tree is grown is considered when corrected class imbalance (referred to as "balanced_subsample"). Little difference in overall performance on the test set was found (Table 5); for convenience, we show the results when using the non-balanced random forest classifier.

Table 5: Performance of various flavours of random forest classifiers on test set

| Flavour of random forest | Macro-F1 on test set |
|---|---|
| Balanced subsample | 97.2% |
| Balanced | 97.3% |
| Not balanced | 97.3% |

### C. Methods of analyzing results

#### *Performance measures*

On an enzymatic dataset, we measure performance for a single enzyme class $i$ through precision ($p_i$) and recall ($r_i$).

$$p_i = \frac{\mathrm{tp_i}}{\mathrm{tp_i} + \mathrm{fp_i}} \quad (17)$$

$$r_i = \frac{\mathrm{tp_i}}{\mathrm{tp_i} + \mathrm{fn_i}} \quad (18)$$

In the above, $\mathrm{tp}_i$, $\mathrm{fp}_i$ and $\mathrm{fn}_i$ represent the number of true positives, false positives and false negatives for the $i^{\mathrm{th}}$ enzyme class, respectively. In terms of summary statistics over all ECs, we macro-average precision and recall so as to equally represent enzyme classes within the dataset, irrespective of class size. We extend the equation for calculating macro-averaged precision and recall as given in [10] as follows:

$$P_M = \frac{\sum_{i=1}^{l_{P\cap R}} p_i}{l_P} \quad (19)$$

$$R_M = \frac{\sum_{i=1}^{l_R} r_i}{l_R} \quad (20)$$

Here, $l_P$ is the number of classes predicted by the method/tool, $l_R$ is the actual number of classes in the dataset, and $l_{P\cap R}$ is the number of classes predicted by the method/tool but also actually present in the dataset. Therefore, (19) penalises the prediction of classes not present in the dataset. As we equally value precision and recall, we often report the performance using the F1-score as given below:

$$F_1 = 2 \times \frac{P_M \times R_M}{P_M + R_M} \quad (21)$$

We note that, unless macro-averaging is specified (such as when annotations between models, or between organisms are compared), we compute precision and recall over the entire dataset as follows (i.e. finding the number of predictions that is either a true positive, false positive or false negative over any class *i*). This is also referred to as micro-averaging [10]:

$$P = \frac{\sum_i \mathrm{tp_i}}{\sum_i \mathrm{tp_i} + \sum_i \mathrm{fp_i}} \quad (22)$$

$$R = \frac{\sum_i \mathrm{tp_i}}{\sum_i \mathrm{tp_i} + \sum_i \mathrm{fn_i}} \quad (23)$$

On a non-enzymatic dataset, we measure performance in terms of specificity (or the true negative rate) as given below.

$$\mathrm{specificity} = \frac{\mathrm{tn}}{\mathrm{tn} + \mathrm{fp}} \quad (24)$$

Here, tn and fp denote the number of true negative and the number of false positives respectively (respectively: the number of non-enzymes correctly not predicted to be enzymes, and incorrectly predicted to be enzymes).

***Performance on multi-functional proteins***

We computed the performance of individual tools and ensemble methods on proteins associated with more than one EC. We identified such proteins as those associated with more than one complete EC number. We used the same measures as given in equations (17)-(21) to compute performance on multifunctional proteins; that is, we treat the predictions of individual ECs (that may or may not co-occur) separately.

When focusing on performance on multi-functional proteins and concerning those ECs that are predictable by all tools, we only consider performance on those multi-functional proteins assigned in SwissProt with all ECs predictable by all tools.

***Performance on test set based on sequence similarity to training data***

We further compared the performance of each classifier on the test dataset, stratified based on its similarity to training data. For these purposes, we employed as measure the maximum test-to-training sequence identity, abbreviated as MTTSI (also used in [1, 11]). For each EC represented in the entire dataset, we calculate the sequence identity of test sequences against training sequences belonging to the same enzyme class (using DIAMOND [12]); then, given a

MTTSI measure of *x%*, we calculate macro-precision and macro-recall for those sequences from the test set sharing a maximum sequence identity of *x%* to any of the corresponding training sequences (restricting ourselves to results with E-value at most 0.1).

### D. Metabolic model reconstruction

#### *KEGG database: curation*

The KEGG database [13] was set up as one of the databases that can be used for metabolic model reconstruction. First, all KEGG reactions and corresponding equations were downloaded. KEGG reaction-EC mappings were used to capture biochemical reactions catalyzed by different enzymes. Post-processing steps were further undertaken to finally produce the reaction database ultimately used for metabolic network reconstruction, comprising 9,509 reactions. These post-processing steps are outlined below.

*Removal of duplicate metabolites and reactions, and macromolecular processing*
KEGG contains instances of equivalent reactions and compounds, especially in the case of glycans which have both glycan identifiers and compound identifiers (respectively of the form Gxxxxx and Cxxxxx, where x is a digit). Such metabolites were identified and a single one retained in the reaction database, duplicate reactions subsequently removed. Reactions involving macromolecular processes were excluded from the reaction database. These were identified by scanning reactions with undefined stoichiometries (such as an *n* instead of an actual number), or with the same compound appearing on both sides of the equation. Reactions marked as incomplete or unclear in KEGG were also removed. Reactions (1,204 in the entire reaction database) involving generic compounds (identified in KEGG as being generic or not having a formula) are separately output for user consideration.

*Determination of reaction reversibility*
Reactions were by default marked as reversible (lower bound of -1000 and upper bound of 1000). However, following the rule of thumb outlined in [14], reactions involving the transfer of a phosphate from an ATP molecule were set as irreversible, with the exception of ATP synthetase (ECs 2.3.3.8 and 6.2.1.18 respectively corresponding to reactions R00352 and R01322).

*Spontaneous and non-enzymatic reactions*
173 biochemical reactions are marked in KEGG as either spontaneous or non-enzymatic (in the "Comment" section of the entry) and are thus considered as being present in network reconstructions irrespective of EC annotation. In the case of KEGG, the following default reactions are also included in all initial draft reconstructions: (i) those enabling the diffusion of small hub metabolites (water, oxygen, carbon dioxide, ammonia, diphosphate, phosphate and protons); (ii) those enabling the interconversion of glucose into its two stereoisomers; (iii) conversion of various fatty acyl-CoA metabolites into a generic acyl-CoA metabolite; and (iv) a reaction accounting for energy expenditure for non-growth associated reasons.

*Balancing reactions*
Reactions were verified for being balanced, including with an R group, by verifying whether the sum of elements on the left and right of the chemical equation match. Those that could be easily

fixed with commonly occurring metabolites were fixed; reactions that remained unbalanced were discarded. In a last step, we verified that reactions were balanced by performing simulations involving the entire reaction database. In short, the reaction database, lacking exchange reactions, represents a closed system through which there can be no metabolite consumption or production; in the presence of imbalance, we expect that there would be a metabolite that could enter or exit the system. For these purposes, we performed flux balance analysis simulations (as described later), setting the objective function as production or consumption of a separate metabolite in each simulation. We then found that there were no metabolites that could either be produced or consumed in these experiments, confirming the nature of our database as a closed system.

#### *BiGG databases and addition of non-EC associated reactions via sequence similarity*

In addition to the KEGG database, we constructed Architect such that it can use various BiGG-based databases for model reconstruction. These are taken from the CarveMe reconstruction tool, and as described in [15], have been curated extensively. Four reaction databases are thus available, in addition to a universal database of metabolism: databases specific to Gram-positive and Gram-negative bacteria, as well as those concerning archaeal and cyanobacterial species. The ECs associated with each gene is parsed out and is used by Architect's model reconstruction module.

Furthermore, given the presence of gene-associated reactions that are not linked to any EC in the BiGG-based databases, Architect uses sequence similarity to find evidence for the presence of such reactions. Therefore, we compiled a database consisting of protein sequences associated with each of these reactions by using information from CarveMe's GitHub repository (https://github.com/cdanielmachado/carveme).

Last, we identified a set of reactions in the BiGG database marked as either spontaneous or non-enzymatic in the different databases (557 and 548 reactions in the archaeal and cyanobacterial databases respectively, and 547 in each of the Gram-positive, Gram-negative and main databases). As in the case of the KEGG reaction database, such reactions are always included in metabolic models reconstructed by Architect.

#### *Relevant simulation details*

Through our gap-filling procedure, we intend to supplement the draft model with reactions necessary for production of biomass (as defined by the user). In particular, we find a small set of reactions whose addition to the model enables flux through the biomass reaction, while prioritizing those with higher confidence from enzyme annotation. The following details mathematical formulations relevant to metabolic simulations using flux balance analysis and related concepts, followed by details of the gap-filling formulation.

*Flux Balance Analysis (FBA)*

Flux Balance Analysis maximizes flux through an objective function (like biomass production) given reaction equations and physicochemical constraints such as lower and upper bounds for reaction fluxes. It is formulated as the following linear programming (LP) problem [16].

$$\max c^T v \tag{25}$$
$$\text{such that } \mathrm{Sv} = 0,$$

$$\text{and } v_L \leq v \leq v_U$$

Given a metabolic model with $m$ metabolites and $n$ reactions, the flux distribution is the only variable in the formulation and is given by $v$, a vector of length $n$ where $v_i$ represents the flux through the $i^{th}$ reaction; $v_L$ and $v_U$ (both vectors of length $n$) constrain the reaction fluxes. $S$ is an $m$ by $n$ stoichiometric matrix, representing the stoichiometries of the $m$ involved metabolites in $n$ model reactions; the constraint "$Sv = 0$" enforces steady state within the system. The $n$-vector $c$ is used to indicate the ratio of reaction fluxes to maximize; in particular, we maximize flux through the objective function (at index $k$) by setting $c_i = 1$ when $i = k$, and $c_i = 0$ otherwise.

*Flux Variability Analysis (FVA)*

For a given fraction ($f$) of the optimal value of the objective function as computed using FBA (α), the allowable range of flux $[v_{i,min}, v_{i,max}]$ through the $i^{th}$ reaction can be computed through flux variability analysis through separate minimization and maximization LP formulations [17]:

$$\begin{gathered} \min/\max v_i \\ \text{such that } \mathrm{Sv} = 0, \\ v_L \leq v \leq v_U, \\ \text{and } c^T \mathrm{v} \geq f \cdot \alpha \end{gathered} \tag{26}$$

*Dead-end metabolites and their identification*

Given the steady-state assumption inherent within FBA, a metabolite which is neither exported nor imported but strictly present within the network can neither accumulate nor deplete; that is, any metabolite produced must be completely consumed, and vice-versa. As a consequence, reactions involved with a metabolite that is either not consumed or not produced (called a dead-end) cannot carry flux [18]. We adopt the following procedure to identify those metabolites that are dead-ends due to network structure and reaction reversibility. We scan each row of the stoichiometric matrix and identify those reactions that either consume or produce the metabolite in question. If the metabolite is involved in only 1 reaction, it is marked as a dead-end. Otherwise, if it is only produced or consumed (the reactions involved also being irreversible), the metabolite is identified as a dead-end.

***Identification of gap-filling candidates required for biomass production***

Given a high-confidence metabolic model and a user-specified biomass reaction (as described in the main text), reactions from the reaction database and exchange reactions for deadend metabolites may still not suffice to produce some biomass components; therefore, we initially find biomass components for which a demand reaction is clearly required. To do so, we first create a universal metabolic network consisting of high-confidence reactions and all other reactions from the entire reaction database; we call this network N. We verify that reactions in the universal network suffice to produce biomass. If not, we perform multiple flux balance analysis (FBA) simulations, individually maximizing production of each biomass component; for each component that cannot be produced, a demand reaction is created. Such a reaction is marked as essential and added to N.

We then prioritize amongst gap-filling candidates as follows. We perform flux variability analysis on gap-filling reactions in N and find those reactions that can carry non-zero flux to produce at least 50% of the optimal flux through the biomass function in the universal database (that is, those having $v_{i,min} \neq 0$ or $v_{i,max} \neq 0$). These constitute our gap-filling candidates. We further mark as essential those gap-filling candidates required for biomass production: such reactions can only carry non-zero flux (that is, $v_{i,min} > 0$ or $v_{i,max} < 0$). The other reactions form our set R of gap-filling candidates.

***Essentiality experiments***

In a reaction knock-out experiment, FBA is performed following the deletion of reactions of interest (setting $v_i = 0$ for the $i^{th}$ reaction). In the case where the value of the objective function is consequently zero (taken here as any value less than 0.0001), the deleted reaction is identified as essential.

In this study, gene essentiality experiments were performed. For gold-standard and CarveMe models, reactions that ought to be deleted following deletion of a gene were found by interpreting Boolean statements representing gene-protein-reaction associations [14] given in the model. If biomass could not be produced, the gene in question was marked as essential. In the case of the models reconstructed by Architect, a different procedure was applied for identifying essential genes given that gene-protein-relationships are not directly predicted by our pipeline. Here, we assume a one-to-one relationship between gene and protein, and an OR-relationship between multiple proteins that may be associated with a particular reaction. We note that as a consequence of the latter assumption, only genes uniquely associated to at least one reaction may be found essential in a model reconstructed by Architect.

### E. Model reconstructions: technical details

Supplementary Table 2 summarizes the information that went into metabolic model reconstruction and accompanying comparisons for the organisms of interest in this paper. The following gives technical details on how the model reconstructions were made using CarveMe, PRIAM and Architect.

*CarveMe reconstructions*

All CarveMe reconstructions were performed as follows:

```
carve <fasta_file>
```

In the case of *E. coli*, we note that running CarveMe with or without minimal media (as defined by CarveMe) yields the same network:

```
carve <fasta_file> -g M9
```

Given that we ran CarveMe without defined media, we only include an import reaction for glucose for the Architect models. To ensure comparability, we used the same biomass as used by CarveMe for model reconstruction in Architect, as indicated in the table above.

*PRIAM reconstructions*

In addition to the specification of the `--complete_genome`, `--cn` and `--nc` flags, all PRIAM-based reconstructions were performing using the following parameters:

```
--pt: 0.5
--mp: 60
--cc: T
```

*Architect reconstructions*

When Architect was run on the BiGG reaction database, non-EC gene-associated reactions were included in the models when predicted at an E-value lower than $10^{-20}$.

When running Architect, the default integrality constraint is set at $10^{-8}$, and can be increased in case of error (such as due to timeout or out-of-memory). This was done in the case of *C. elegans*, whose reconstruction with the BiGG database was performed under a reduced integrality constraint of $10^{-7}$.

Links to external databases are included in the SBML output using the MetaNetX database [19].

### F. Additional results

#### 3.1a Filtering multifunctional enzyme predictions has minimal impact on performance

Architect's enzyme annotation module, as presented in the text, reports all high-confidence predictions made for each protein thus offering the possibility of annotating a protein with multiple enzyme activities. Such functionality is of interest given that approximately 5% of enzymes annotated by SwissProt are multifunctional (**Supplemental Figure 1C**). Consequently, such enzyme annotation tools as PRIAM (Claudel-Renard, Chevalet et al. 2003), EFICAz (Kumar and Skolnick 2012) and EnzDP (Nguyen, Srihari et al. 2015) have additional logic for filtering multiple EC-annotations. Here, we investigated integrating the following heuristic with the random forest, logistic regression and naïve Bayes classifiers: proteins are assigned the highest-scoring EC and allowed additional high-confidence ECs only if they co-occur with the top-scoring EC at least 10 times in the training data. Comparing the performance of the ensemble methods on subsections of the data composed only of single- and multifunctional proteins, we found that the application of the heuristic improved macro-precision for the single-functional proteins, but with a significant decrease in the macro-recall of the multifunctional proteins (**Supplemental Figures 4 and 5**). Given that the addition of this heuristic impacts performance on this class of proteins, Architect's enzyme annotation module outputs all EC predictions.

#### 3.4 Comparing gene essentiality predictions

Beyond enzyme annotations, we were interested in comparing the performance of models generated by Architect and CarveMe in metabolic flux-based simulations exploring predictions of gene essentiality. PRIAM-based reconstructions were excluded from these comparisons as they require additional refinements to be used as models of metabolic flux. Further, only models based on the two bacterial species (*N. meningitidis* and *E. coli*) were examined to avoid the potentially confounding influence of assigning reactions to specific subcellular compartments. In the subsequent comparisons, precision and recall were computed with reference to gene deletion studies performed *in vivo* [20, 21]. In general, there is little difference in the precision and recall of CarveMe and Architect (whether using the KEGG or the BiGG database for model reconstruction or relying only on EC annotations obtained from either EnzDP or PRIAM; **Figure 3 and Supplemental Figures 12 and 13**). However, lower recall was obtained when DETECT predictions were used in isolation, highlighting again the value of high predictive range in tools involved in model reconstruction. At the same time, models generated from all reconstruction tools generally exhibited low recall with respect to predicting gene essentiality. This relatively high rate of false negatives may be explained by several factors including: (1) certain essential genes may have been excluded from reconstructed models or misassigned function; (2) key Boolean relationships between multiple genes associated with a single reaction — such as with heteromeric enzyme complexes [14] — may not be captured in the models; or (3) the biomass equation used during model simulations may be incomplete. Interestingly, of the genes experimentally found to be essential, 78% and 83% were incorporated into Architect models for *N. meningitidis* and *E. coli* respectively built using KEGG; however, most of these (43% and 37% respectively) were not predicted to be essential (**Supplemental Table 1**), suggesting avenues for improving Architect by either limiting pathways predicted from ECs (thereby reducing pathway redundancy and highlighting the essentiality of certain genes), or through better representations of gene-protein-reaction relationships.

### G. Supplementary Tables

Supplementary Table 1: Overlap of *in silico* determined essential genes with those found essential *in vivo*.

| | Method | # TP | # FP | # FN (altogether) | # essential genes not included in output model | Precision | Recall | Recall (only considering genes included in model) |
|---|---|---|---|---|---|---|---|---|
| *N. meningitidis* | Architect-KEGG | 80 | 23 | 160 | 53 | 77.7% | 33.3% | 42.8% |
| | Architect-BiGG | 89 | 18 | 151 | 73 | 83.2% | 37.1% | 53.3% |
| | CarveMe | 84 | 21 | 156 | 45 | 80.0% | 35.0% | 43.1% |
| | Arch-DETECT | 47 | 17 | 193 | 74 | 73.4% | 19.6% | 28.3% |
| | Arch-EnzDP | 77 | 26 | 163 | 72 | 74.8% | 32.1% | 45.8% |
| | Arch-PRIAM | 77 | 20 | 163 | 60 | 79.4% | 32.1% | 42.8% |
| *E. coli* | Architect-KEGG | 76 | 14 | 173 | 43 | 84.4% | 30.5% | 36.9% |
| | Architect-BiGG | 51 | 15 | 198 | 61 | 77.3% | 20.5% | 27.1% |
| | CarveMe | 57 | 8 | 192 | 39 | 87.7% | 22.9% | 27.1% |
| | Arch-DETECT | 37 | 15 | 212 | 81 | 71.2% | 14.9% | 22.0% |
| | Arch-EnzDP | 64 | 15 | 185 | 59 | 81.0% | 25.7% | 33.7% |
| | Arch-PRIAM | 53 | 15 | 196 | 47 | 77.9% | 21.3% | 26.2% |

Supplementary Table 2: Comparisons of various aspects of model reconstruction for *C. elegans*, *N. meningitidis* and *E. coli*. The number of reactions in reconstructed models that are not blocked (and corresponding number of metabolites) is indicated within brackets, except in the case of automatically reconstructed *C. elegans* models (*). The number of exchange reactions added by Architect for deadend metabolites is given within brackets (**).

| | | Organism | | |
|---|---|---|---|---|
| | | *C. elegans* | *N. meningitidis* | *E. coli* |
| Architect, CarveMe and PRIAM-based reconstructions | Source of protein sequences | WormBase database | Ensembl database | UniProt Proteome ID: UP000000625 |
| | Num of protein sequences | 20,483 | 2,063 | 4,391 |
| Architect reconstruction | Biomass used | Main CarveMe biomass (using KEGG identifiers when using KEGG database; same as for other organisms) | Gram-negative CarveMe biomass | Gram-negative CarveMe biomass |
| Architect reconstruction using KEGG | Num of protein sequences | 1,389 | 387 | 967 |
| | Num of reactions* | 1,433 | 885 (347) | 1,671 (864) |
| | Num of metabolites* | 1,530 | 1,082 (297) | 1,689 (605) |
| | Num of gap-filling reactions** | 32 (14) | 20 (14) | 10 (7) |
| Architect reconstruction using KEGG and predictions from individual tools | Num of reactions* | DETECT: 1,025<br>EnzDP: 1,271<br>PRIAM: 1,356 | DETECT: 773 (301)<br>EnzDP: 814 (323)<br>PRIAM: 861 (342) | DETECT: 1,108 (449)<br>EnzDP: 1,500 (712)<br>PRIAM: 1,697 (852) |
| | Num of metabolites | DETECT: 1,221<br>EnzDP: 1,402<br>PRIAM: 1,474 | DETECT: 960 (256)<br>EnzDP: 1,013 (282)<br>PRIAM: 1,060 (295) | DETECT: 1,329 (350)<br>EnzDP: 1,586 (505)<br>PRIAM: 1,720 (595) |
| | Num of gap-filling reactions** | DETECT: 38 (27)<br>EnzDP: 42 (26)<br>PRIAM: 41 (25) | DETECT: 44 (31)<br>EnzDP: 31 (25)<br>PRIAM: 27 (21) | DETECT: 44 (30)<br>EnzDP: 23 (18)<br>PRIAM: 19 (17) |
| Architect reconstruction using BiGG | Num of protein sequences | 432 | 298 | 670 |
| | Num of reactions* | 1,409 | 1,600 (527) | 3,023 (2,356) |
| | Num of metabolites* | 1,312 | 1,513 (400) | 2,049 (1,387) |
| | Num of gap-filling reactions** | 25 (10) | 29 (12) | 5 (2) |
| CarveMe reconstruction | Biomass used | Main CarveMe biomass | Gram-negative biomass | Gram-negative biomass |
| | Num of genes/protein sequences | 561 | 613 | 1,639 |
| | Num of reactions* | 1,538 | 1,569 (1,541) | 2,833 (2,810) |
| | Num of metabolites* | 1,100 | 1,138 (1,110) | 1,735 (1,717) |

| | | | | |
|---|---|---|---|---|
| Reconstruction using PRIAM | Num of protein sequences | 920 | 452 | 1,052 |
| | Num of reactions | 1,387 | 867 | 1,890 |
| | Num of metabolites | 1,397 | 935 | 1,717 |
| Gold-standard models | Provenance | WormJam [22]; version 2019_01_01 from *https://gh.wormjam.life* | Nmb_iTM560 [21] | iML1515 [20] |
| | Num of genes | 1,520 | 559 | 1,515 |
| | Num of reactions* | 3,632 (2,947) | 1,527 (not available) | 2,719 (2,459) |
| | Num of metabolites* | 2,833 | 1,297 | 1,192 |
| UniProt gold-standard annotations | Provenance | Uniprot | Uniprot | Using annotations for UP000000625 in SwissProt |
| | Num of protein sequences with EC annotations when comparing against KEGG-based Architect | 1,446 | 495 | 1,123 |
| | Num of protein sequences with EC annotations when comparing against BiGG-based Architect | 659 | 504 | 1,111 |
| Essentiality results | Provenance | Not applicable | [21] | [20] |

Supplementary Table 3: Breakdown of annotations of SwissProt sequences by individual and ensemble methods into true positives, true negatives, false positives and false negatives

| | | Enzymatic test set | | | Non-enzymatic test set | | | |
|---|---|---|---|---|---|---|---|---|
| | | # TPs | # FPs | # FNs | # prots annotated as enzymes | # FP annotations | # TN proteins | Sum of FP annotations |
| Individual tools | CatFam | 31,364 | 3,644 | 13,209 | 5,090 | 5,159 | 288,977 | 8,803 |
| | DETECT_all | 34,525 | 19,762 | 10,048 | 16,262 | 24,544 | 277,805 | 44,306 |
| | DETECT_high | 34,158 | 1,476 | 10,415 | 2,110 | 2,163 | 291,957 | 3,639 |
| | EFICAz_all | 32,466 | 5,099 | 12,107 | 3,909 | 4,283 | 290,158 | 9,382 |
| | EFICAz_high | 31,606 | 2,569 | 12,967 | 1,497 | 1,505 | 292,570 | 4,074 |
| | EnzDP_all | 39,649 | 134,150 | 4,924 | 49,229 | 151,923 | 244,838 | 286,073 |
| | EnzDP_high | 37,930 | 821 | 6,643 | 1,587 | 1,760 | 292,480 | 2,581 |
| | PRIAM_all | 43,099 | 2,430 | 1,474 | 4,293 | 4,870 | 289,774 | 7,300 |
| | PRIAM_high | 42,845 | 1,227 | 1,728 | 2,149 | 2,597 | 291,918 | 3,824 |
| Ensemble methods | Majority rule | 42,030 | 1,362 | 2,543 | 7,763 | 8,501 | 286,304 | 9,863 |
| | EC-specific tool | 43,430 | 479 | 1,143 | 10,500 | 10,961 | 283,567 | 11,440 |
| | Naïve Bayes | 43,404 | 281 | 1,169 | 2,308 | 2,418 | 291,759 | 2,699 |
| | L1-regression | 43,559 | 291 | 1,014 | 8,983 | 9,248 | 285,084 | 9,539 |
| | L2-regression | 43,567 | 290 | 1,006 | 8,815 | 9,084 | 285,252 | 9,374 |
| | Random forest | 43,595 | 297 | 978 | 8,601 | 8,899 | 285,466 | 9,196 |

Supplementary Table 4: Breakdown of organism-specific annotations by ensemble and individual tool into true positive, false positive and false negative. This comparison is done against UniProt annotations for sequences used in model reconstruction, and the predictions from the ensemble method come the naïve Bayes classifier for ECs in Architect's training database, and PRIAM (high-confidence) otherwise.

| | Tool | # TP | # FP in all | # FP on enzymes | # FP on non-enzymes (# proteins in brackets) | # FP on proteins with partial_ECs | # FN |
|---|---|---|---|---|---|---|---|
| *C. elegans* | DETECT | 825 | 753 | 113 | 495 (486) | 145 | 763 |
| | EnzDP | 995 | 545 | 83 | 390 (335) | 72 | 593 |
| | PRIAM | 1,170 | 667 | 194 | 413 (374) | 60 | 418 |
| | Architect | 1,260 | 867 | 117 | 648 (613) | 102 | 328 |
| *N. meningitidis* | DETECT | 355 | 109 | 45 | 44 (43) | 20 | 206 |
| | EnzDP | 415 | 75 | 28 | 28 (26) | 19 | 146 |
| | PRIAM | 460 | 96 | 41 | 32 (29) | 23 | 101 |
| | Architect | 482 | 108 | 42 | 44 (41) | 22 | 79 |
| *E. coli* | DETECT | 648 | 231 | 144 | 29 (28) | 58 | 762 |
| | EnzDP | 1,087 | 186 | 120 | 33 (26) | 33 | 323 |
| | PRIAM | 1,253 | 145 | 92 | 16 (15) | 37 | 157 |
| | Architect | 1,257 | 173 | 102 | 24 (23) | 47 | 153 |
